# gReLU: A comprehensive framework for DNA sequence modeling and design

**DOI:** 10.1101/2024.09.18.613778

**Authors:** Avantika Lal, Laura Gunsalus, Surag Nair, Tommaso Biancalani, Gokcen Eraslan

## Abstract

Deep learning models are increasingly being used to perform a variety of tasks on DNA sequences, such as predicting tissue- and cell type-specific sequence activity, deriving cis-regulatory rules, predicting non-coding variant effects, and designing synthetic regulatory sequences. However, these models require specialized knowledge to build, train and interpret correctly. In addition, the field is hampered by the lack of interoperability between models and software built by different groups. Here we present gReLU, a comprehensive software framework that enables users to easily execute advanced sequence modeling pipelines, including data preprocessing, model training, hyperparameter tuning, evaluation, interpretation, variant effect prediction, and design of novel regulatory elements. The software is accompanied by a model zoo containing state-of-the-art pre-trained models that can easily be downloaded, applied, and fine-tuned. This framework and resources will accelerate research across the field of DNA sequence modeling and enable the effective design of synthetic regulatory elements.

## 1. Introduction

Deep learning models are increasingly used as powerful oracles in DNA sequence analysis. By training on DNA sequences coupled with data from diverse functional genomic assays, these models learn the cis-regulatory code in different biological contexts and can make predictions on unseen sequences (Eraslan et al. 2019; Lal 2022). Thus, they can be queried to perform *in silico* experiments, such as predicting the effect of sequence variants or synthetic DNA constructs. Such sequence models have enabled prioritization of disease-associated non-coding variants (Trevino et al. 2021), *in silico* genome engineering (Martyn et al. 2023), and design of cell type specific regulatory elements (Taskiran et al. 2024; Gosai et al. 2023). Further, interpreting the regulatory rules learned by the models has the potential to reveal novel cis-regulatory mechanisms (Avsec, Weilert, et al. 2021).

However, these models are difficult to build, train and interpret correctly, and minor errors can easily result in misleading predictions (Whalen et al. 2022). In addition, the field is hampered by the lack of interoperability between models and software. Different groups have built models using different frameworks, such as Keras (Avsec, Weilert, et al. 2021; Avsec, Agarwal, et al. 2021; Linder et al. 2023) and PyTorch (Chen et al. 2019, 2022; Lal et al. 2021). Instead of a common underlying framework being adapted to create new models, it is common to see a new model architecture accompanied by unique custom code to load and process data and evaluate and interpret the model. Thus it can be difficult to combine and compare models built by different groups, or to fine-tune models for novel tasks. Further, many individual tools have been built to handle specific modeling tasks, such as sequence design (Gosai et al. 2023; Schreiber and Lu 2020) or model interpretation (Shrikumar et al. 2018), making it difficult to chain these tasks into a workflow.

Previous groups have highlighted this issue and have attempted to build unifying software frameworks (Chen et al. 2019; Avsec et al. 2019), including a recently developed python package for predictive modeling based on PyTorch (Klie et al. 2023). However, these frameworks are still limited in scope, and there remains a need for comprehensive tools that encompass multiple workflows and keep pace with recent advances in the field, such as novel architectures (Linder et al. 2023; Avsec, Agarwal, et al. 2021), loss functions (Avsec, Weilert, et al. 2021; Linder et al. 2023), genomic foundation models (Dalla-Torre et al. 2023; Nguyen et al. 2023), training models on single-cell datasets (Yuan and Kelley 2023), simulation-based interpretation methods (Avsec, Weilert, et al. 2021; Toneyan and Koo 2024), and model-guided sequence design algorithms (Schreiber and Lu 2020; Linder and Seelig 2021). In particular, there remains a disconnect between the software ecosystems surrounding different types of models, such as large-scale models that predict many outputs over long genomic contexts (Avsec, Agarwal, et al. 2021; Linder et al. 2023), smaller task- or locus-specific models (Avsec, Weilert, et al. 2021), and models trained on single-cell data (Hepkema et al. 2023; Virshup et al. 2023).

Here, we present gReLU, a python-based software framework that comprehensively integrates and enables sequence modeling pipelines, including data loading, preprocessing, training, testing, fine-tuning, interpretation, simulation, variant effect prediction, and sequence design. Through example analyses, we demonstrate how gReLU enables multiple state-of-the-art sequence modeling workflows on diverse types of datasets and models. Further, we release a zoo of large pre-trained models that can easily be downloaded, used, and fine-tuned using our software. We believe that these contributions have the potential to dramatically accelerate the field of DNA sequence modeling.

## 2. Results

### 2.1 Overview of gReLU

gReLU takes as input DNA sequences or genomic coordinates along with functional data, which can be presented in a variety of common genomic formats. It enables the user to perform preprocessing tasks, such as filtering the data and splitting it into training, validation and test sets; to construct neural networks with provided architectures or custom architectures; and to train these models to predict functional information from DNA sequence. Using their trained models or models downloaded from the accompanying model zoo, users can benchmark models on held-out data, predict the functions of novel sequences, predict the effects of sequence variants or perturbations, and design DNA sequences with desired activity using the trained model as an oracle **(Fig. 1)**.

**Fig 1:**
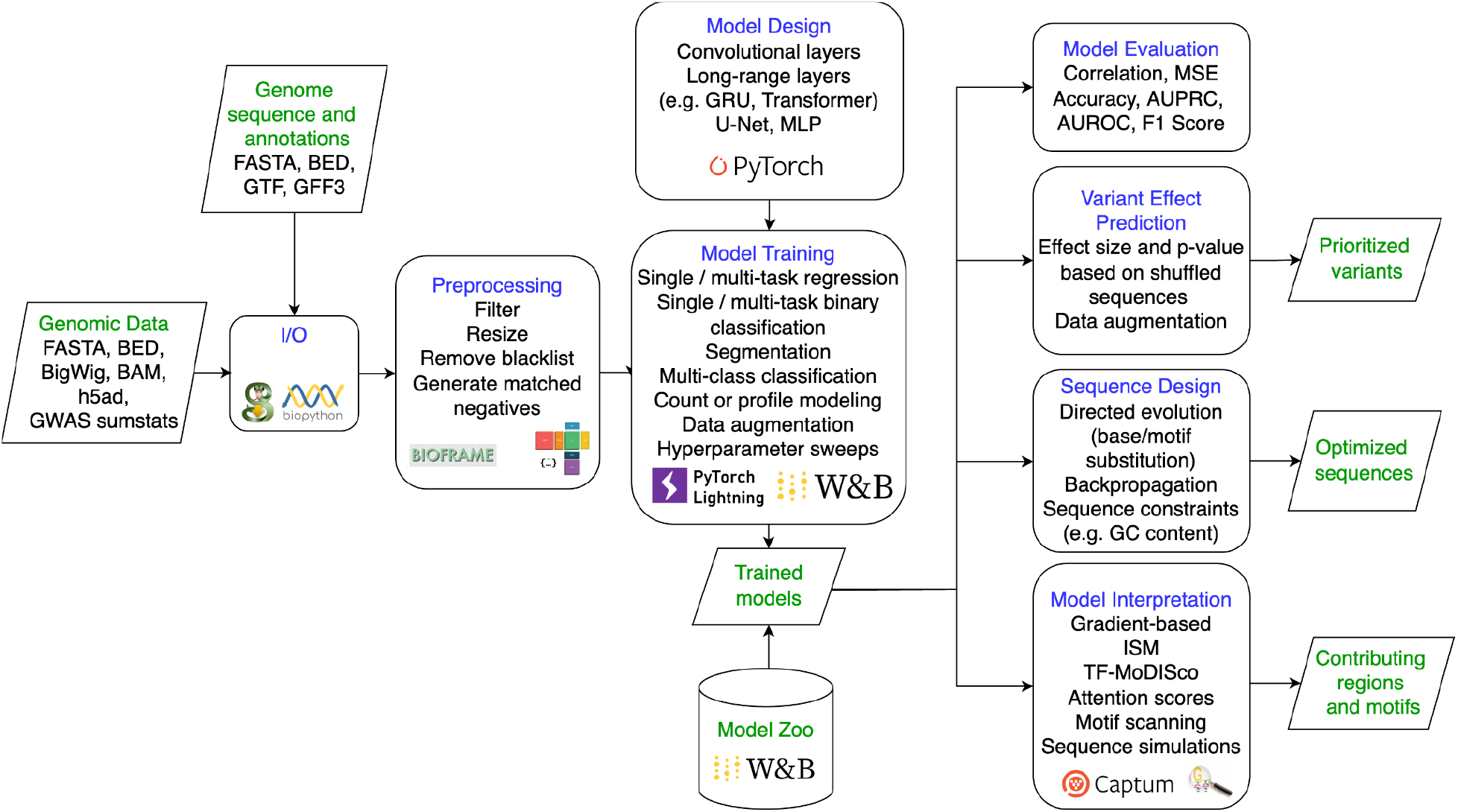
An overview of the gReLU package.

#### Data Input

DNA sequences and associated functional data can be loaded from standard data formats including tabular files, FASTA, BED, BigWig, fragment files and AnnData objects (Virshup et al. 2021). Given a set of genomic coordinates and a reference genome name, we use genomepy (Frölich et al. 2023) to load genome sequences and annotations from genome databases and extract sequences.

#### Data Processing

gReLU has a suite of preprocessing functions including filtering blacklist regions (Amemiya, Kundaje, and Boyle 2019), filtering sequences based on their activity, quality or content, finding GC-matched negative control regions in the genome, and calculating per-base sequencing coverage from fragment files. Genomic regions can be split into sets for training, validation and testing randomly or by chromosome. gReLU has several custom pytorch dataset classes to load genomic data from each input format, convert sequences into one-hot encoded tensors, and feed the resulting sequences and labels to the model in batches. gReLU also provides optional augmentation functions to increase the available data and train more robust models. These include reverse complementation, shifting and jittering the sequence regions used for training, and introducing random mutations into the sequence.

#### Model Design

gReLU provides several model architectures that the user can customize to their needs, ranging from small convolutional models to large transformer-based model architectures such as Enformer (Avsec, Agarwal, et al. 2021) and Borzoi (Linder et al. 2023). Users can also define their own architecture by combining pre-built layers and blocks defined in gReLU, including convolutional, recurrent, transformer, U-Net and dense layers.

#### Model Training

gReLU can train models to perform single-task or multi-task regression, single-task or multi-task binary classification, and multi-class classification of sequences. Suitable loss functions are provided for each task. Specifically, Mean Squared Error (MSE), Poisson and multinomial (Avsec, Weilert, et al. 2021; Linder et al. 2023) losses for regression, binary cross-entropy loss for binary classification and cross-entropy loss for multi-class classification. Users can also assign weights to specific classes or examples in order to focus the model on important or rare sequence categories.

Training is performed using PyTorch Lightning, which enables multi-GPU parallelization. gReLU also optionally enables logging and hyperparameter sweeps using Weights and Biases. Models are evaluated on the validation set at every epoch and the model with the lowest validation loss is saved. To ensure reproducibility, gReLU saves model checkpoints containing not only the weights of the model but also metadata concerning the model architecture and training data.

#### Model Inference and Evaluation

Models can be evaluated on a held-out test set. We offer several performance metrics tailored for different tasks, such as Mean Squared Error (MSE), Pearson correlation, Cross-entropy, Accuracy, Area Under the Precision-Recall Curve (AUPRC), Area Under the Receiver-Operator Characteristic (AUROC), and F1 score. gReLU also checks classification models for miscalibration. These functions enable users to easily benchmark models and compare the performance of different architectures and hyperparameters. For models that predict coverage profiles, gReLU can return predictions along with the coordinates of the corresponding genomic regions, and can provide the input genomic coordinates corresponding to any portion of the model’s output, accounting for cropping and binning of the model’s predictions.

#### Model Interpretation

gReLU provides functions for nucleotide-resolution importance scoring using *in silico* mutagenesis (ISM), DeepSHAP or gradient-based attribution methods. It also runs TF-MoDISco (Shrikumar et al. 2018) on the output in order to identify common motifs and matches them to known motifs using TOMTOM (S. Gupta et al. 2007). Notably, users can score the importance of each input nucleotide to complex functions of the model’s output, such as the ratio between the model’s predictions in two different cell types, or over specific genomic regions.

Sequences can be scanned with user-provided motifs or motifs from the JASPAR database (Castro-Mondragon et al. 2022) using FIMO (Grant, Bailey, and Noble 2011) and this can be used to annotate the regions with high importance scores. gReLU also includes functions to perform simulation experiments in which regions of a sequence are shuffled, or motifs are inserted into random sequences in order to understand the regulatory grammar learned by the model (Avsec, Weilert, et al. 2021).

#### Variant Effect Prediction

gReLU accepts sequence variants, extracts the surrounding genomic sequence for each variant and creates batches of sequences containing the reference and alternate alleles. gReLU can use any trained model to perform inference on the reference- and alternate-allele containing genomic sequences and can compare the results to calculate an effect size for each variant. It can also perform data augmentation and average the variant effect across both DNA strands and shifted sequences to obtain more robust estimates. In addition, gReLU can compute a background distribution for each variant by shuffling the genomic sequences while conserving dinucleotide frequency, before inserting the reference and alternate alleles and computing effect sizes. This allows the computation of p-values along with effect sizes for all variants.

#### Sequence Design

gReLU contains simple and flexible functions that enable model-based sequence design using any trained model as an oracle. It currently includes three design approaches:

##### 1. Directed Evolution by *in silico* mutagenesis

Given a set of initial sequences, all possible single-base substitutions are introduced, the altered sequences are evaluated using the trained model, and the best sequence is chosen based on a user-specified objective function. This is repeated for a user-specified number of iterations.

##### 2. Directed Evolution by motif insertion

Given a set of initial sequences and a set of sequence motifs, each motif is inserted into all possible positions in the input sequences, the resulting sequences are evaluated using the trained model, and the best sequence is chosen based on a user-specified objective function. This is repeated for a user-specified number of iterations.

##### 3. Gradient-based optimization while optionally penalizing edits (Ledidi)

gReLU integrates the Ledidi method (Schreiber and Lu 2020) which uses gradient descent to iteratively update an initial sequence in order to achieve a user-specified objective. Optionally, the user can specify a penalty on the number of edits introduced into the initial sequence, encouraging the exploration of local sequence space.

Users can define the objective to optimize using these methods, which can range from directly increasing or decreasing the model’s prediction or complex functions of the model’s output - for example, increasing or decreasing the ratio between the model’s predictions in two different cell types, or over specific genomic regions. Users can also specify a subset of sequence positions to be mutated. In directed evolution, users can also introduce penalties to encourage or discourage specific sequence patterns, such as motifs, GC content or CpG instances in the optimized sequences.

#### Model zoo

gReLU is accompanied by a model zoo, which is hosted on Weights and Biases. gReLU provides users with functions to programmatically search the model zoo and download any model. The model zoo stores not only the weights and hyperparameters for each model but also details of the training data and the preprocessing steps used to create it. The zoo currently includes the following models:

1. Enformer (Avsec, Agarwal, et al. 2021), including both mouse and human heads;
2. Borzoi (Linder et al. 2023), including 4 replicate models with both mouse and human heads;
3. A binary classification model trained on CATlas snATAC-seq data from human tissues (Lal et al. 2023; Zhang et al. 2021);
4. A multiclass classification model trained on the universal chromatin state annotations for the human genome (Lal et al. 2024; Vu and Ernst 2022);
5. A regression model trained on DNase-seq data from GM12878 cells, discussed below.

Below, we present two experiments demonstrating the use of gReLU for biologically relevant applications.

### 2.2 Modeling chromatin accessibility and predicting non-coding variant effects

Deciphering the mechanisms through which non-coding variants alter gene regulation and ultimately cellular state is crucial for understanding disease biology. DNA sequence models have shown promise in predicting the effect of variants on transcription factor binding, chromatin accessibility, and gene expression across diverse cellular contexts (Zhou and Troyanskaya 2015; Eraslan et al. 2019; Trevino et al. 2021). To illustrate how gReLU can be used to nominate regulatory variants, we trained a model using gReLU to predict DNase-seq signal in the GM12878 cell line (ENCODE Project Consortium 2012). The model takes as input 2114 bp of sequence, and outputs the log of the total DNase-seq counts in the central 1000 bp of the input sequence. We used a dilated convolutional architecture, similar to BPNet (Avsec, Weilert, et al. 2021), consisting of 9 convolution layers with 256 channels each. The model was trained on DNase-seq peaks as well as a GC-content matched set of non-peak regions chosen by gReLU. The model was trained for a total of 50 epochs, and the best model based on validation set Pearson correlation was chosen. The Pearson correlation between measured and predicted DNase-seq signal on sequences in the test set was 0.80 **(Fig. 2A)**.

**Fig 2:**
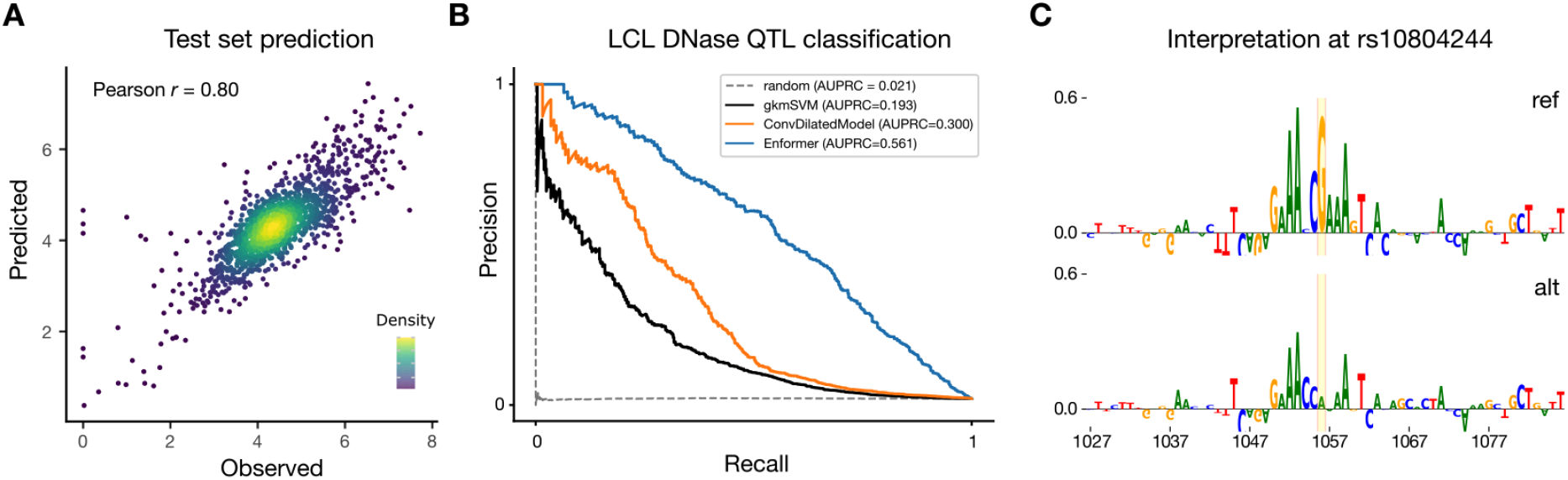
A) Test set predictions for a convolutional regression model trained on GM12878 DNase-seq using gReLU **B)** Precision-recall curve for scoring 28,309 single-nucleotide variants out of which 574 are DNase-seq quantitative trait loci (dsQTLs) in lymphoblastoid cell lines using the regression model and other baselines **C)** Nucleotide-resolution importance scores obtained from the regression model for the reference and alternate alleles of rs10804244 using the input x gradient method highlight a disrupted IRF motif. The variant is highlighted in yellow.

To assess the model’s ability to identify variants that modulate chromatin accessibility, we computed predicted variant effects for a set of 28,309 single-nucleotide variants, of which ∼2% are dsQTLs identified in a collection of lymphoblastoid cell lines (LCLs) (Degner et al. 2012; Lee et al. 2015). The regression model achieved an area under precision recall curve (AUPRC) of 0.3 at classifying dsQTLs in this dataset, outperforming a random predictor, as well as a previously published gkmSVM model (Lee et al. 2015) (Fig 2B).

Leveraging gReLU’s model zoo, we easily benchmarked the Enformer model (Avsec, Weilert, et al. 2021) on the same dataset **(Fig. 2B)**. Enformer’s higher AUPRC of 0.561 likely reflects the advantages conferred from its long input sequence length, profile modeling, and multi-species training.

Finally, we used gReLU to interpret the predictions of the regression model at rs10804244 by scoring the importance of each surrounding nucleotide with the input x gradient method and scanning the sequence with TF binding motifs. Together, the results suggest that the G>A variant weakens a binding site for the IRF transcription factor **(Fig 2C)**.

### 2.3 *In silico* cellular engineering using long-context models

We applied the Borzoi model (Linder et al. 2023) from the public gReLU model zoo to demonstrate the editing capabilities of the gReLU package. Our goal was to edit a native enhancer to alter the expression of its target gene in a tissue-specific manner (**Fig. 3A**). We selected the *PPIF* gene, which is expressed across various tissues but shows particularly strong expression in blood and comparatively lower expression in other tissues, such as the brain.

**Figure 3:**
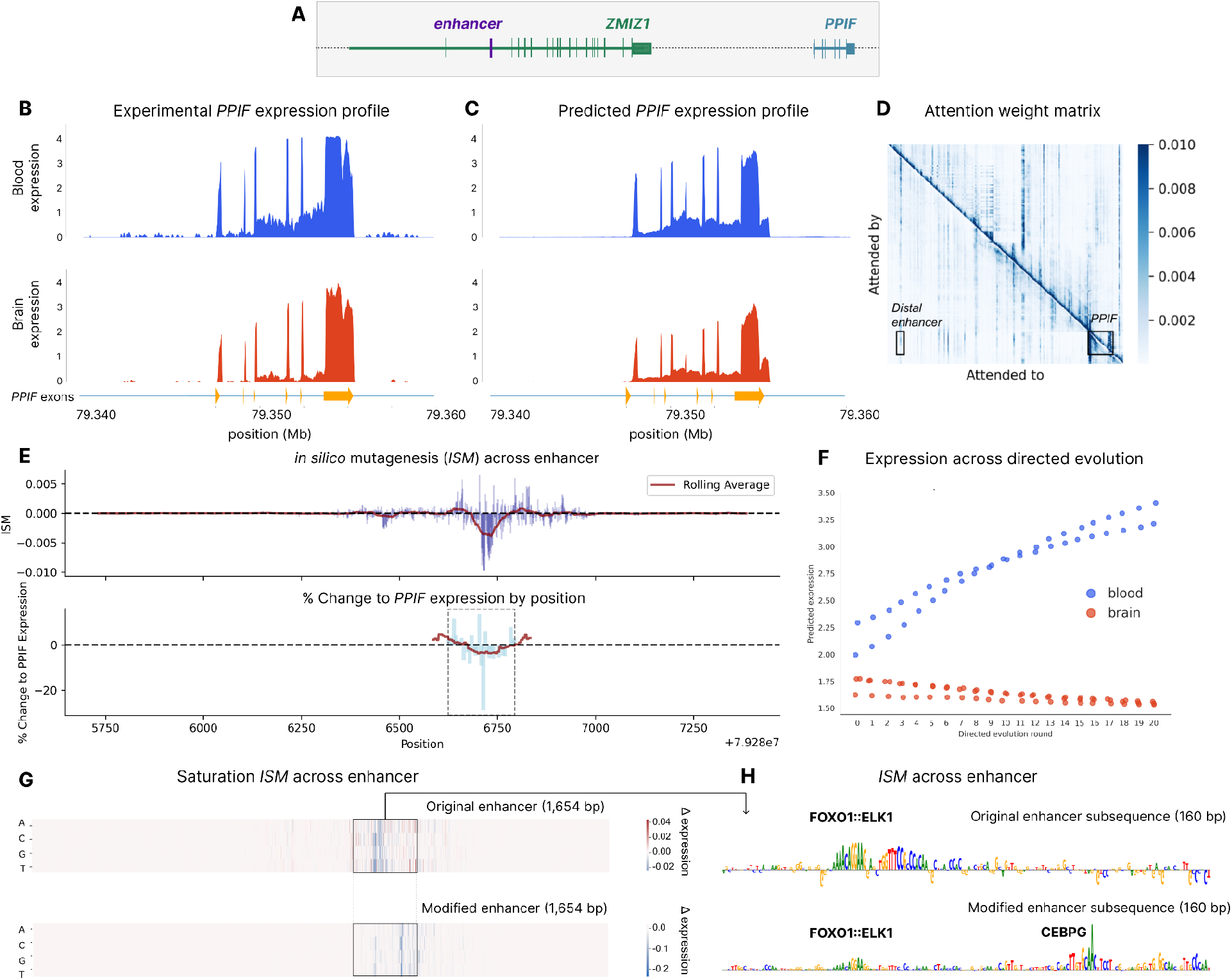
In silico modifications of a distal enhancer impacting PPIF expression. **A)** Schematic representation of the PPIF locus, highlighting the enhancer region modified at chr10:79285732-79287386. **B)** Mean expression profile of PPIF across two GTEx whole blood tracks (blue) and three GTEx brain tracks (red). **C)** Borzoi-predicted mean PPIF expression profile for the same six GTEx tracks. **D)** Attention weight matrix averaged across all heads of the final transformer layer of Borzoi shows increased attention between the PPIF gene and the distal PPIF enhancer. **E)** Functional analysis of the distal enhancer. Mean saturation in silico mutagenesis (ISM) across the region, with a rolling average shown in red (top). Mean change in PPIF expression induced by 5-bp edits, as measured by Variant FlowFISH in K562 cells (bottom, Martyn et al.) **F)** Changes in predicted blood and brain expression over the course of 20 rounds of directed evolution across the distal enhancer, optimized to maximize the difference in expression between the two tissues. Points represent the predicted RNA-seq coverage over exons of the PPIF gene in two GTEx tracks in blood and three GTEx tracks in brain. **G)** Comparative ISM tracks illustrating the changes in predicted PPIF expression induced by single base pair edits across the original enhancer sequence (top) and edited enhancer sequence (bottom). **H)** Detailed ISM tracks of a subsequence within the enhancer, demonstrating the emergence of novel motifs in the edited blood-specific enhancer.

Using gReLU, we visualized the predicted RNA-seq coverage profile of the *PPIF* gene and found that it closely mirrored the ground-truth expression data from GTEx (**Fig. 3B, C**).

The Borzoi model has a long input context spanning 524,288 bases, enabling the model to integrate information from distal regulatory elements. Using gReLU’s interpretation functions, we visualized the attention matrix from the model’s final attention layer, allowing us to examine distal sequence elements that the model attends to when predicting RNA-seq coverage over the *PPIF* gene (**Fig. 3D**). These included a previously characterized enhancer located 61.7kb upstream of the transcription start site (TSS) (Martyn et al. 2023).

gReLU’s interpretation functions further allow users to estimate the contribution of each input nucleotide to defined functions of the output, such as coverage over specific genomic regions and/or in a subset of output tasks. To investigate the regulatory activity of this enhancer, we performed *in silico* mutagenesis (ISM) across the enhancer sequence, specifically quantifying the impact of base changes on whole blood expression across *PPIF* exons. Our analysis revealed that a central location is particularly sensitive to mutagenesis (**Fig. 3E, top**). Our results mirrored experimental findings (Martyn et al. 2023). Mutations at the center of the sequence resulted in a loss in *PPIF* expression upon perturbation, while mutations surrounding the center resulted in a modest increase in expression (**Fig. 3E, bottom**).

We next sought to edit this enhancer to increase the tissue bias of *PPIF* expression. We defined our objective as the difference between the average blood expression across the PPIF exons and the average brain expression across the PPIF exons. Next, we applied directed evolution using gReLU to make 20 individual base pair edits with the goal of increasing this metric. This iterative mutagenesis resulted in a 54.26% increase in blood expression and a 10.26% decrease in brain expression (**Fig. 3F**). ISM across the original and modified enhancers points to the same sensitivity at the center of the sequence (**Fig. 3G**). Using gReLU’s motif scanning functionality (Rauluseviciute et al. 2024), we observed that our edits created a novel CEBPG motif, in addition to the FOXO1:ELK1 motif present in the original sequence (**Fig. 3H**). The CEBP family is involved in hematopoiesis and myeloid cell differentiation (Avellino and Delwel 2017).

In sum, we demonstrate how gReLU may be used to investigate genomic loci using publicly available models, interpret them through *in silico* mutagenesis and motif identification, and edit them to achieve specific tissue expression profiles.

## 3. Discussion

Several efforts have been made to address the fragmentation of computational tools in the sequence modeling field (Chen et al. 2019; Klie et al. 2023; Avsec et al. 2019), demonstrating the importance of this problem. gReLU aims to enhance this software ecosystem by enabling users to easily and reliably apply, fine-tune and interpret numerous state-of-the-art models such as Borzoi (Linder et al. 2023), to process data and train custom models employing recently developed state-of-the-art architectures, loss functions and interpretation methods, and to design novel regulatory elements by optimizing several properties simultaneously. To our knowledge, this combination of functionalities is currently not available in any existing package.

Potential future extensions of gReLU include incorporating additional types of sequence models into the same framework, such as masked language models (Nguyen et al. 2023; Dalla-Torre et al. 2023), or models based on state-of-the-art efficient layers that can model long genomic contexts (Poli et al. 2023; Gu and Dao 2023), as well as incorporating additional sequence design algorithms (Gosai et al. 2023; Vaishnav et al. 2022), enabling efficient training on larger multi-species datasets, modeling sequencing biases in genomic assays, and modeling individual genomes (Drusinsky, Whalen, and Pollard 2024).

## 4. Methods

### 4.1 Processing of GM12878 DNase-seq data

From the ENCODE portal, we downloaded data corresponding to the GM12878 DNase-seq experiment ENCSR000EJD. Specifically, we downloaded the read-depth normalized bigwig (ENCFF093VXI) and the peaks narrowPeak file (ENCFF588OCA), aligned to the hg38 reference genome. Using gReLU, we extended the peaks 250bp on each side of the summit and merged overlapping regions. We filtered the regions to autosomes and removed regions overlapping blacklisted regions (Amemiya, Kundaje, and Boyle 2019). We obtained a set of GC-matched negative regions using the grelu.data.preprocess.get_gc_matched_intervals function with binwidth=0.05. We used sequences from Chromosome 10 as the validation set and sequences from Chromosome 11 as the test set.

### 4.2 Architecture and training of the DNase-seq regression model

We used the DilatedConvModel architecture provided in gReLU with the following parameters: channels=256, n_conv=9, input length 2114 bp and output lengths of 1000 bp. We created a dataset using the BigWigSeqDataset class with label_aggfunc=“sum”, label_transform_func=np.log1p, rc=True, max_seq_shift=3, max_pair_shift=50, augment_mode=“random”. Thus, the labels are transformed to the log of the summed counts over the output 1000 bp region. We trained the model using the mean squared error (MSE) loss. We used a learning rate of 10^−4^, batch size 128, and trained for a total of 50 epochs, keeping track of the model with the highest Pearson correlation on the validation set.

### 4.3 Variant Effect Prediction and Interpretation

We downloaded the previously curated list of 574 LCL dsQTLs SNPs and 27,735 control SNPs (Lee et al. 2015). We scored the variants using grelu.variant.predict_variant_effects with compare_func=“subtract” and genome=“hg19”, using the provided hg19 coordinates. This returns the log fold-change (LFC) of the predicted counts between the reference and the alternate alleles. We assessed performance using the sklearn.metrics.average_precision_score function with the provided labels and the absolute value of the predicted LFC. We generated the model interpretations using the grelu.interpret.score.get_attributions function with method=“inputxgradient” and visualized them with the grelu.visualize.plot_attributions function.

### 4.4 Inference and interpretation using the Borzoi model

The Borzoi model was downloaded from the public gReLU model zoo. Matched ground-truth RNA-seq data was acquired from the Borzoi training dataset. We selected blood due to its high PPIF expression. It is also the closest GTEx track matching K562 and CD8+ lymphocyte models used in Martyn et al. The tracks presented in Fig. 3B and C represent the averaged expression across the two GTEx blood RNA-seq tracks and three GTEx brain RNA-seq tracks (dataset IDs: GTEX-1LB8K-0005-SM-DIPED.1, GTEX-1OKEX-0006-SM-DKPQ2.1, GTEX-13FTY-0011-R11a-SM-5IJEA.1, GTEX-1EX96-0011-R4a-SM-ARU82.1, GTEX-1H3O1-1726-SM-9WYSR.1).

The results shown in Figure 3 were obtained using Borzoi replicate 0. ISM was computed with respect to the average expression from the two GTEx blood tracks across all bins overlapping the *PPIF* exons. We present the log2 ratio between the predictions for the mutated sequence and the reference sequence. Similarly, predicted expression values in Fig. 3F were calculated as the average prediction across all bins overlapping the *PPIF* exons. Motif scanning was performed using the JASPAR 2024 database using core non-redundant motifs in vertebrates.

### 4.5 Experimental evidence

We conducted *in silico* saturation mutagenesis on a *PPIF* regulatory region located at chr10:79285732-79287386 (hg38). To validate our computational results, we compared them with experimentally validated insertion experiments at the same enhancer region performed by Martyn et al (Martyn et al. 2023). In these experiments, 5bp windows were replaced across tiles of the locus. The results from the PPIF Enhancer Tiling Mutagenesis were obtained from Supplementary Table 8, and the coordinates were converted from hg19 to hg38 using LiftOver (Nassar et al. 2023). For comparison, we calculated the mean ISM results per base. We then applied a rolling average with a window size of 50bp for the ISM results and 80bp for the experimental results.

## 5. Code Availability

gReLU is available under an MIT license at https://github.com/Genentech/gReLU.

Documentation is available at https://genentech.github.io/gReLU/.

Tutorials are available at https://github.com/Genentech/gReLU/tree/main/docs/tutorials.

Code used to perform the analyses in this paper is available at https://github.com/Genentech/gReLU-applications.

The model zoo is available at https://wandb.ai/grelu/ and includes the models described in this paper.

## 7. Author Contributions

A.L. and G.E. built the package. L.G. and S.N. performed analyses. T.B. provided mentorship and supervision. A.L., G.E., L.G., and S.N. wrote the manuscript. All authors read and approved the manuscript.

## 8. Acknowledgments

We would like to thank David Garfield, Oriol Fornes, Kipper Fletez-Brant, Muhammed Hasan Celik, Carlo de Donno, Jayaram Kancherla, Jean-Philippe Fortin, and Jorge Kageyama for their advice, feedback and contributions to the package.

## References

Amemiya, Haley M., Anshul Kundaje, and Alan P. Boyle. 2019. “The ENCODE Blacklist: Identification of Problematic Regions of the Genome.” Scientific Reports 9 (1): 9354.

Avellino, Roberto, and Ruud Delwel. 2017. “Expression and Regulation of C/EBPα in Normal Myelopoiesis and in Malignant Transformation.” Blood 129 (15): 2083–91.

Avsec, Žiga, Vikram Agarwal, Daniel Visentin, Joseph R. Ledsam, Agnieszka Grabska-Barwinska, Kyle R. Taylor, Yannis Assael, John Jumper, Pushmeet Kohli, and David R. Kelley. 2021. “Effective Gene Expression Prediction from Sequence by Integrating Long-Range Interactions.” Nature Methods 18 (10): 1196–1203.

Avsec, Žiga, Roman Kreuzhuber, Johnny Israeli, Nancy Xu, Jun Cheng, Avanti Shrikumar, Abhimanyu Banerjee, et al. 2019. “The Kipoi Repository Accelerates Community Exchange and Reuse of Predictive Models for Genomics.” Nature Biotechnology 37 (6): 592–600.

Avsec, Žiga, Melanie Weilert, Avanti Shrikumar, Sabrina Krueger, Amr Alexandari, Khyati Dalal, Robin Fropf, et al. 2021. “Base-Resolution Models of Transcription-Factor Binding Reveal Soft Motif Syntax.” Nature Genetics 53 (3): 354–66.

Castro-Mondragon, Jaime A., Rafael Riudavets-Puig, Ieva Rauluseviciute, Roza Berhanu Lemma, Laura Turchi, Romain Blanc-Mathieu, Jeremy Lucas, et al. 2022. “JASPAR 2022: The 9th Release of the Open-Access Database of Transcription Factor Binding Profiles.” Nucleic Acids Research 50 (D1): D165–73.

Chen, Kathleen M., Evan M. Cofer, Jian Zhou, and Olga G. Troyanskaya. 2019. “Selene: A PyTorch-Based Deep Learning Library for Sequence Data.” Nature Methods 16 (4): 315–18.

Chen, Kathleen M., Aaron K. Wong, Olga G. Troyanskaya, and Jian Zhou. 2022. “A Sequence-Based Global Map of Regulatory Activity for Deciphering Human Genetics.” Nature Genetics 54 (7): 940–49.

Dalla-Torre, Hugo, Liam Gonzalez, Javier Mendoza-Revilla, Nicolas Lopez Carranza, Adam Henryk Grzywaczewski, Francesco Oteri, Christian Dallago, et al. 2023. “The Nucleotide Transformer: Building and Evaluating Robust Foundation Models for Human Genomics.” bioRxiv. 10.1101/2023.01.11.523679.

Degner, Jacob F., Athma A. Pai, Roger Pique-Regi, Jean-Baptiste Veyrieras, Daniel J. Gaffney, Joseph K. Pickrell, Sherryl De Leon, et al. 2012. “DNase I Sensitivity QTLs Are a Major Determinant of Human Expression Variation.” Nature 482 (7385): 390–94.

Drusinsky, Shiron, Sean Whalen, and Katherine S. Pollard. 2024. “Deep-Learning Prediction of Gene Expression from Personal Genomes.” bioRxiv. 10.1101/2024.07.27.605449.

ENCODE Project Consortium. 2012. “An Integrated Encyclopedia of DNA Elements in the Human Genome.” Nature 489 (7414): 57–74.

Eraslan, Gökcen, Žiga Avsec, Julien Gagneur, and Fabian J. Theis. 2019. “Deep Learning: New Computational Modelling Techniques for Genomics.” Nature Reviews. Genetics 20 (7): 389–403.

Frölich, Siebren, Maarten van der Sande, Tilman Schäfers, and Simon J. van Heeringen. 2023. “Genomepy: Genes and Genomes at Your Fingertips.” Bioinformatics 39 (3). 10.1093/bioinformatics/btad119.

Gosai, S. J., R. I. Castro, N. Fuentes, J. C. Butts, S. Kales, R. R. Noche, K. Mouri, P. C. Sabeti, S. K. Reilly, and R. Tewhey. 2023. “Machine-Guided Design of Synthetic Cell Type-Specific Cis -Regulatory Elements.” bioRxiv : The Preprint Server for Biology, August. 10.1101/2023.08.08.552077.

Grant, Charles E., Timothy L. Bailey, and William Stafford Noble. 2011. “FIMO: Scanning for Occurrences of a given Motif.” Bioinformatics 27 (7): 1017–18.

Gu, Albert, and Tri Dao. 2023. “Mamba: Linear-Time Sequence Modeling with Selective State Spaces.” arXiv [cs.LG]. arXiv. http://arxiv.org/abs/2312.00752.

Gupta, Shobhit, John A. Stamatoyannopoulos, Timothy L. Bailey, and William Stafford Noble. 2007. “Quantifying Similarity between Motifs.” Genome Biology 8 (2): R24.

Hepkema, Jacob, Nicholas Keone Lee, Benjamin J. Stewart, Siwat Ruangroengkulrith, Varodom Charoensawan, Menna R. Clatworthy, and Martin Hemberg. 2023. “Predicting the Impact of Sequence Motifs on Gene Regulation Using Single-Cell Data.” Genome Biology 24 (1): 189.

Klie, Adam, David Laub, James V. Talwar, Hayden Stites, Tobias Jores, Joe J. Solvason, Emma K. Farley, and Hannah Carter. 2023. “Predictive Analyses of Regulatory Sequences with EUGENe.” Nature Computational Science 3 (11): 946–56.

Lal, Avantika. 2022. “Deciphering the Regulatory Syntax of Genomic DNA with Deep Learning.” Journal of Biosciences 47. https://www.ncbi.nlm.nih.gov/pubmed/36222139.

Lal, Avantika, Laura Gunsalus, Anay Gupta, Tommaso Biancalani, and Gokcen Eraslan. 2023. “Polygraph: A Software Framework for the Systematic Assessment of Synthetic Regulatory DNA Elements.” bioRxiv. 10.1101/2023.11.27.568764.

Lal, Avantika, Zachary D. Chiang, Nikolai Yakovenko, Fabiana M. Duarte, Johnny Israeli, and Jason D. Buenrostro. 2021. “Deep Learning-Based Enhancement of Epigenomics Data with AtacWorks.” Nature Communications 12 (1): 1507.

Lal, Avantika, David Garfield, Tommaso Biancalani, and Gokcen Eraslan. 2024. “regLM: Designing Realistic Regulatory DNA with Autoregressive Language Models.” bioRxiv. 10.1101/2024.02.14.580373.

Lee, Dongwon, David U. Gorkin, Maggie Baker, Benjamin J. Strober, Alessandro L. Asoni, Andrew S. McCallion, and Michael A. Beer. 2015. “A Method to Predict the Impact of Regulatory Variants from DNA Sequence.” Nature Genetics 47 (8): 955–61.

Linder, Johannes, and Georg Seelig. 2021. “Fast Activation Maximization for Molecular Sequence Design.” BMC Bioinformatics 22 (1): 510.

Linder, Johannes, Divyanshi Srivastava, Han Yuan, Vikram Agarwal, and David R. Kelley. 2023. “Predicting RNA-Seq Coverage from DNA Sequence as a Unifying Model of Gene Regulation.” bioRxiv. 10.1101/2023.08.30.555582.

Martyn, Gabriella E., Michael T. Montgomery, Hank Jones, Katherine Guo, Benjamin R. Doughty, Johannes Linder, Ziwei Chen, et al. 2023. “Rewriting Regulatory DNA to Dissect and Reprogram Gene Expression.” bioRxiv : The Preprint Server for Biology, December. 10.1101/2023.12.20.572268.

Nassar, Luis R., Galt P. Barber, Anna Benet-Pagès, Jonathan Casper, Hiram Clawson, Mark Diekhans, Clay Fischer, et al. 2023. “The UCSC Genome Browser Database: 2023 Update.” Nucleic Acids Research 51 (D1): D1188–95.

Nguyen, Eric, Michael Poli, Marjan Faizi, Armin Thomas, Callum Birch-Sykes, Michael Wornow, Aman Patel, et al. 2023. “HyenaDNA: Long-Range Genomic Sequence Modeling at Single Nucleotide Resolution.” ArXiv, June. https://www.ncbi.nlm.nih.gov/pubmed/37426456.

Poli, Michael, Stefano Massaroli, Eric Nguyen, Daniel Y. Fu, Tri Dao, Stephen Baccus, Yoshua Bengio, Stefano Ermon, and Christopher Ré. 2023. “Hyena Hierarchy: Towards Larger Convolutional Language Models.” arXiv [cs.LG]. arXiv. http://arxiv.org/abs/2302.10866.

Rauluseviciute, Ieva, Rafael Riudavets-Puig, Romain Blanc-Mathieu, Jaime A. Castro-Mondragon, Katalin Ferenc, Vipin Kumar, Roza Berhanu Lemma, et al. 2024. “JASPAR 2024: 20th Anniversary of the Open-Access Database of Transcription Factor Binding Profiles.” Nucleic Acids Research 52 (D1): D174–82.

Schreiber, Jacob, and Yang Young Lu. 2020. “Ledidi: Designing Genomic Edits That Induce Functional Activity.” bioRxiv. 10.1101/2020.05.21.109686.

Shrikumar, Avanti, Katherine Tian, Žiga Avsec, Anna Shcherbina, Abhimanyu Banerjee, Mahfuza Sharmin, Surag Nair, and Anshul Kundaje. 2018. “Technical Note on Transcription Factor Motif Discovery from Importance Scores (TF-MoDISco) Version 0.5.6.5.” arXiv [cs.LG]. arXiv. http://arxiv.org/abs/1811.00416.

Taskiran, Ibrahim I., Katina I. Spanier, Hannah Dickmänken, Niklas Kempynck, Alexandra Pančíková, Eren Can Ekşi, Gert Hulselmans, et al. 2024. “Cell-Type-Directed Design of Synthetic Enhancers.” Nature 626 (7997): 212–20.

Toneyan, Shushan, and Peter K. Koo. 2024. “Interpreting Cis-Regulatory Interactions from Large-Scale Deep Neural Networks for Genomics.” bioRxiv : The Preprint Server for Biology, March. 10.1101/2023.07.03.547592.

Trevino, Alexandro E., Fabian Müller, Jimena Andersen, Laksshman Sundaram, Arwa Kathiria, Anna Shcherbina, Kyle Farh, et al. 2021. “Chromatin and Gene-Regulatory Dynamics of the Developing Human Cerebral Cortex at Single-Cell Resolution.” Cell 184 (19): 5053–69.e23.

Vaishnav, Eeshit Dhaval, Carl G. de Boer, Jennifer Molinet, Moran Yassour, Lin Fan, Xian Adiconis, Dawn A. Thompson, Joshua Z. Levin, Francisco A. Cubillos, and Aviv Regev. 2022. “The Evolution, Evolvability and Engineering of Gene Regulatory DNA.” Nature 603 (7901): 455–63.

Virshup, Isaac, Danila Bredikhin, Lukas Heumos, Giovanni Palla, Gregor Sturm, Adam Gayoso, Ilia Kats, et al. 2023. “The Scverse Project Provides a Computational Ecosystem for Single-Cell Omics Data Analysis.” Nature Biotechnology 41 (5): 604–6.

Virshup, Isaac, Sergei Rybakov, Fabian J. Theis, Philipp Angerer, and F. Alexander Wolf. 2021. “Anndata: Annotated Data.” bioRxiv. 10.1101/2021.12.16.473007.

Vu, Ha, and Jason Ernst. 2022. “Universal Annotation of the Human Genome through Integration of over a Thousand Epigenomic Datasets.” Genome Biology 23 (1): 9.

Whalen, Sean, Jacob Schreiber, William S. Noble, and Katherine S. Pollard. 2022. “Navigating the Pitfalls of Applying Machine Learning in Genomics.” Nature Reviews. Genetics 23 (3): 169–81.

Yuan, Han, and David R. Kelley. 2023. “Author Correction: scBasset: Sequence-Based Modeling of Single-Cell ATAC-Seq Using Convolutional Neural Networks.” Nature Methods 20 (1): 162.

Zhang, Kai, James D. Hocker, Michael Miller, Xiaomeng Hou, Joshua Chiou, Olivier B. Poirion, Yunjiang Qiu, et al. 2021. “A Single-Cell Atlas of Chromatin Accessibility in the Human Genome.” Cell 184 (24): 5985–6001.e19.

Zhou, Jian, and Olga G. Troyanskaya. 2015. “Predicting Effects of Noncoding Variants with Deep Learning-Based Sequence Model.” Nature Methods 12 (10): 931–34.

